# SWEET11 and 15 as key players in seed filling in rice

**DOI:** 10.1101/198325

**Authors:** Jungil Yang, Dangping Luo, Bing Yang, Wolf B. Frommer, Joon-Seob Eom

## Abstract

- Despite the relevance of seed filling mechanisms for crop yield, we still have only a rudimentary understanding of the pathways and transport processes for supplying the caryopsis with sugars. We hypothesized that the recently identified SWEET sucrose transporters may play important roles in nutrient import pathways in the rice caryopsis.
- We used a combination of mRNA quantification, histochemical analyses, translational promoter-reporter fusions and analysis of *knock out* mutants created by genomic editing to evaluate the contribution of SWEET transporters to seed filling.
- In rice caryopses, *SWEET11* and *15* had the highest mRNA levels and proteins localized to four key sites: the nucellus proper at early stages, the nucellar projection close to the dorsal vein, the nucellar epidermis that surrounds the endosperm, and the aleurone. *ossweet11*;*15* double *knock-out* lines accumulated starch in the pericarp while caryopses did not contain a functional endosperm.
- Jointly, SWEET11 and 15 show all hallmarks of being responsible for seed filling with sucrose efflux function at the nucellar projection and transfer across the nucellar epidermis/aleurone interface, delineating two major steps for apoplasmic seed filling, observations that are discussed in relation to observations made in rice and barely on the relative prevalence of these two potential import routes.

## Introduction

Population growth is expected to lead to an increasing need for rice production, especially in Africa (Sharma, 2014). A key question is thus how we can obtain maximal yield potential. Rice grains are composed mainly of starch (over 90% in many cases; www.knowledgebank.irri.org/ricebreedingcourse/Grain_quality.htm), which derives from imported soluble carbohydrates. These carbohydrates are produced in the leaves with the help of photosynthesis and are exported mainly as sucrose via the phloem, which contains ~600 mM sucrose (Fukumorita & Chino, 1982). In order to fill the seeds, and more specifically to generate starch, cell walls and provide energy, sucrose has to be imported into developing caryopses. Phloem strands enter the seed coat, where sucrose is unloaded and transferred into the developing caryopsis to supply cells with nutrients, in particular sugars as sources of energy and as carbon skeletons for cell wall and starch biosynthesis.

In plants, cell-to-cell transport of sugars is thought to be mediated by apoplasmic (export from one cell by a plasma membrane transporter and subsequent import into the adjacent cell by another transport protein) or by symplasmic transfer via plasmodesmata. Other routes are conceivable, but evidence for vesicular transport processes are sparse (van den Broek *et al.*, 1997). Two of the key processes for long distance translocation are phloem loading and seed filling. The transport processes that ultimately lead to cell wall synthesis and storage product accumulation in seeds, in particular in cereal caryopses are not fully understood. To delineate sym- and apoplasmic pathways in rice caryopses, Oparka carried out a combination of ultrastructural and dye tracer studies in the early 80s (Oparka & Gates, 1981a,b, 1982, 1984). The rice caryopsis is supplied by three vascular bundles that pass through the pericarp. The dorsal vascular bundle is the major route for sugar delivery to the developing caryopsis (Oparka & Gates, 1981a; Krishnan & Dayanandan, 2003). He found symplasmic connections between the parenchyma of the dorsal vascular bundle and the nucellar projection as well as a lipid barrier between the inner integument and the nucellar epidermis that blocks the apoplasmic route, therefore requiring transporters for uptake and release at this site for a transcellular pathway. From these comprehensive analyses that included radiotracer analyses using ^14^CO_2_, he proposed that after sucrose had been unloaded from the phloem, it diffuses along the nucellar epidermis from where an unknown set of sugar transporters transfers the sugar into the developing endosperm. Since sucrose, upon arrival in the caryopsis through the phloem of the dorsal vascular bundle, is partially hydrolyzed by cell wall invertases (cwINVs; in particular OsGIF1/OsCIN2) into glucose and fructose, one would predict the presence of both sucrose and hexose transport pathways (Wang *et al.*, 2008). At present, two classes of plasma membrane sucrose transporters (SUTs and clade 3 SWEETs) and three classes of plasma membrane hexose transporters (MSTs (STPs), ERDs and clade 1 and 2 SWEETs (Chen *et al.*, 2015a) are known. SWEETs are a class of seven transmembrane hexose and sucrose uniporters that function as oligomers (Xuan *et al.*, 2013). Their roles in Arabidopsis include nectar secretion, seed filling, and they have been shown to act as susceptibility factors for pathogen infections. It has been proposed that AtSWEET13 functions as a ' revolving door' mechanism to accelerate the transport efficacy (Feng & Frommer, 2015; Latorraca *et al.*, 2017; Han *et al.*, 2017)

In legume seeds, during early developmental stages, cwINVs produce hexoses from the incoming sucrose. These hexoses are thought to stimulate mitotic activity to increase cell number, while subsequently at later stages the cwINVs are switched off and sucrose transporters are induced. Sucrose is thought to then act as a differentiation signal that triggers storage product accumulation (Weber *et al.*, 2005). Specialized sucrose facilitators (SUFs, members of the SUT family), PsSUF1, PsSUF4 and PvSUF1 are appeared sucrose efflux in seed coats of pea and common bean (Zhou *et al.*, 2007). In monocots, specifically in maize and rice, cwINVs also play important roles in seed filling (Cheng & Chourey, 1999; Wang *et al.*, 2008). Several hexose transporters that are likely involved in import of the cwINV-derived hexoses into the caryopsis or endosperm have been identified. Rice *MST4* is expressed during grain developmental stages in maternal tissues including the dorsal vascular bundle, nucellus including nucellar projection and nucellar epidermis, and aleurone layer of the filial endosperm (Wang *et al.*, 2007). *ZmSWEET4c* in maize is expressed in the BETL and necessary for seed filling and BETL differentiation.

The apparent ortholog *OsSWEET4* also appears to have a role in grain filling, although its detailed cell specificity and expression profile has yet to be determined (Sosso *et al.*, 2015). The rice sucrose transporters *OsSUT1*, *OsSUT3* and *OsSUT4* localize to the aleurone (Ishimaru *et al.*, 2001; Bai *et al.*, 2016). Antisense inhibition of *OsSUT1* expression caused seed filling defects (Scofield *et al.*, 2002). Since we found that several sucrose transporting SWEETs contribute to seed filling in Arabidopsis (Chen *et al.*, 2015b), we speculate one or several orthologs in rice may play analogous roles in supplying sucrose to either the rice SUTs or cwINVs.

Here, we show that similar as in legumes, the hexose transporter *OsSWEET4* is predominantly expressed during early stages of caryopsis development, while OsSWEET11 and 15 mRNAs accumulated at higher levels during later developmental stages. Expression was found in the ovular vascular trace, nuclear epidermis and endosperm. We also demonstrate that *ossweet11* single mutant and *ossweet11;15* double mutants show retarded endosperm development and defected endosperm filling. Another work performed others in parallel also identified OsSWEET11 as key to seed filling in rice (Ma *et al.*, 2017). The phenotype of the *ossweet11;15* double mutants was more severe in having an empty seed phenotype, while the pericarp of *osweet11* and *osweet11;15* mutants accumulated more starch. These results indicate that OsSWEET11 and OsSWEET15 are necessary for sugar efflux from the maternal nuclear epidermis as well as efflux from the ovular vascular trace to the apoplasm and may also contribute to sucrose influx into the aleurone.

## Materials and Methods

### Plant materials and growth conditions

The *Oryza sativa* ssp. *japonica* cultivar Kitaake was used for CRISPR-Cas9 and TALEN mediated mutagenesis in *OsSWEET11* and *OsSWEET15* genes. The methods for CRISPR-Cas9 induced mutant line (*ossweet11-1*) was described previously (Zhou *et al.*, 2014). Briefly, guide RNA genes targeting the start codon ATG containing sequence (5’-TCACCAGTAGCAATGGCAGG-3’) of *OsSWEET11* was used with Cas9 for transformation (see Fig. 2S). While method for TALEN-induced mutant lines (*ossweet11-2*, *ossweet15-1* and *ossweet15-2*) was also described (Li *et al.*, 2014). One pair of TALENs for *OsSWEET11* and another pair of TALENs for *OsSWEET15* were designed and engineered for rice transformation. For detailed sequence and location of TALEN targets, see Fig. 2S. Double mutants (*ossweet11-1; 15-1 and ossweet11-2; 15-2*) were created by crossing. Rice wild-type plants and mutants were grown either in field conditions (Stanford University Campus, CA, USA) or in greenhouses (Stanford and ISU) under long-day conditions (14h day/10h night, 28-30°C).

### Genotyping of rice plants

Rice genomic DNA was extracted using CTAB method. PCR was performed using ExTaq DNA polymerase (Clontech, Mountain View, CA, USA) with a melting temperature of 56°C and 53°C for *OsSWEET11* and *OsSWEET15,* respectively (for primers, see Supplementary Table 1). The PCR-amplicons from the mutant alleles has been sequenced. Chromatograms were read and aligned using BioEdit.

### RNA isolation and transcript analyses

Total RNA was isolated using the SpectrumTM Plant total RNA kit (Sigma, St. Louis, MO, USA) or the Trizol method (Invitrogen, Carlsbad, CA, USA) and first strand cDNA was synthesized using Quantitect reverse transcription Kit (Qiagen, Hilden, Germany). qRTPCR to determine expression level was performed using the LightCycler 480 system (Roche, Penzberg, Germany), with the 2^-Δ*Ct*^ method for relative quantification. Specific primers for *OsSWEET11* and *OsSWEET15* were used and *OsUBI1* was served as internal control gene (see Supplementary Table 1).

## Generation of *OsSWEET11* and *OsSWEET15* reporter constructs

The 2,334 bp GUSplus coding sequence with nopaline synthase terminator were amplified by PCR from pC1305.1 (Cambia, Canberra and Brisbane, Australia). The amplified fragment was subcloned into the pJET2.1/blunt vector (Thermo Fisher, Waltham, MA, USA) and confirmed by sequencing. The cloned fragment digested with *Sac*I and *EcoR*I was further subcloned into the plant transformation vector pC1300intC to generate a promoterless GUSplus coding vector. For tissue specificity analysis, total 4,354 bp genomic clone containing 2,106 bp of the 5' upstream region and 2,248 bp of the entire coding region of the *OsSWEET11* gene and total 4,193 bp genomic clone containing 2,069 bp of the 5’ upstream region and 2,124 bp of the entire coding region of the *OsSWEET15* gene without stop codon was amplified by PCR using Kitaake genomic DNA as a template, respectively (primers: Supplementary Table 1). The amplified product was sub-cloned into a pJET2.1/blunt vector and confirmed by sequencing. The cloned fragment digested with *Hind*III and *BamH*I for *OsSWEET11* and *Xba*I and *Kpn*I for *OsSWEET15* were further inserted in front of GUSplus coding sequence digested with *Hind*III/*BamH*I and *Xba*I/*Kpn*I, respectively. The resulting *pOsSWEET11*:gOsSWEET11-GUSplus and *pOsSWEET15*:gOsSWEET15-GUSplus constructs were used to transform rice Kitaake. 18 and 13 independent lines were obtained for *pOsSWEET11*:gOsSWEET11-GUSplus and *pOsSWEET15*:gOsSWEET15-GUSplus, respectively, with similar expression pattern.

### Histochemical GUS analyses

Rice immature seeds at 5 DAP (days after pollination) from *pOsSWEET11*:gOsSWEET11-GUS-8 and *pOsSWEET15*:gOsSWEET15-GUS-14 were collected and stained to test cell-type specific expression pattern analysis in developing caryopses. Samples were collected in 90% cold acetone for fixation, vacuum infiltrated for 10 min and incubated for 30 min at room temperature. Seeds were vacuum infiltrated in staining buffer [Staining solution w/o 5-bromo-4-chloro-3-indole-beta-glucuronide (XGluc)] on ice for 10 min. Solution was changed with GUS staining solution [50 mM sodium phosphate (pH 7.0), 10 mM EDTA, 20% (v/v) methanol, 0.1 % (v/v) Triton X-100, 1mM potassium ferrocyanide, 1mM potassium ferricyanide, 2 mM 5-bromo-4-chloro-3-indole-beta-glucuronide (X-Glc) dissolved in dimethyl sulfoxide]. Samples were incubated at 37°C. After 20 min staining, samples were incubated series of ethanol (20%, 35%, 50%) at room temperature for 30 min each. For paraffin section, seeds were fixed using FAA for 30 min [50% (v/v) ethanol, 3.7% (v/v) formaldehyde, 5% (v/v) acetic acid]. Dehydration was performed with ethanol series (70%, 80%, 90%, 100%, each for 30 min) and 100 % tert-butanol. Samples were transferred and embedded in Histosec pastilles (Millipore, Billerica, MA, USA). Cross sections (8 μm) were performed by rotary microtome (Jung RM 2025, Wetzlar, Germany). Specimens were observed with Nikon eclipse e600 microscope. Images for GUS histochemistry in Figure 1 were enhanced uniformly in Photoshop by adjusting brightness (+5), contrast (+7), and blue color balance (+4) to increase the ability to observe the X-gluc staining.

### Plastic embedding and sectioning

Wild-type, *ossweet11-1* and *ossweet11-1;15-1* seeds were fixed using 4% PFA [1X PBS buffer (37 mM NaCl, 10 mM Na_2_HPO_4_, 2.7 mM KCl, 1.8mM KH_2_PO_4_ and a pH of 7.4) with 4% paraformaldehyde], vacuum infiltrated for 15 min and incubated overnight. Dehydration of samples were performed in ethanol with a series of concentration (10%, 30%, 50%, 75% and 95%). Plastic embedding was performed accordingly to the LR White embedding kit protocol (Electron Microscopy Science). Cross-sections (1μ m) were performed using Ultracut (Reichert-Jung, Wetzlar, Germany), stained with 0.1% Safranin O for 30 second and washed twice with distilled water, followed by 3 min starch staining for 3 min with Lugol’ s staining solution. Specimens were observed with Nikon eclipse e600 microscope.

### FRET sucrose sensor analysis in HEK293T cells

The *OsSWEET11* and *OsSWEET15* coding sequences were cloned into the Gateway entry vector pDONR221f1, and then cloned into vector pcDNA3.2V5 by LR recombination reaction for expression in HEK293T cells. HEK293T cells were co-transfected with a plasmid carrying the *OsSWEET11* or *OsSWEET15* and the sucrose sensor FLIPsuc90μ-sCsA, using Lipofectamine 2000 (Invitrogen, Carlsbad, CA, USA). For FRET imaging, HBSS medium was used to perfuse HEK293T/ FLIPsuc90μ-sCsA cells pulsed with 20 mM sucrose. Image acquisition and analysis were performed as previously described (Chen *et al.*, 2012). Empty vector was served as a negative control.

## Results

### Identification of OsSWEET11 and OsSWEET15 in rice caryopses

To identify SWEETs that are expressed specifically in rice caryopses, we analyzed public microarray data from RiceXPro (ricexpro.dna.affrc.go.jp). We focused on members of clade 3 since they had been shown to function as plasma membrane sucrose transporters. Among the five clade 3 SWEETs analyzed, *OsSWEET11* and *15* had the highest mRNA levels in the endosperm between 7 and 14 DAP (Fig. S1). To validate the micro-array data, we harvested immature seeds at different developmental stages from greenhouse-grown plants and re-analyzed mRNA levels of *OsSWEET11* and *15* by qRT-PCR. For comparison, we analyzed *OsSWEET4*, which had been shown to play an important role as a hexose transporter in seed development (Sosso *et al.*, 2015)(Fig. 1). At 1 DAP, *SWEET4* had the highest mRNA levels, but already at 3 DAP, levels had declined about ~3-fold. In comparison, *OsSWEET11* was low at DAP1, but equal to *OsSWEET4* at 3 DAP. *OsSWEET11* gradually increased throughout seed development. While in the microarrays *OsSWEET15* was only 2-3x lower compared to *OsSWEET11*, our qRT-PCR analysis indicated a much lower relative level however the developmental pattern of *OsSWEET15* mRNA levels was similar to that of *OsSWEET11*.

**Fig. 1.**
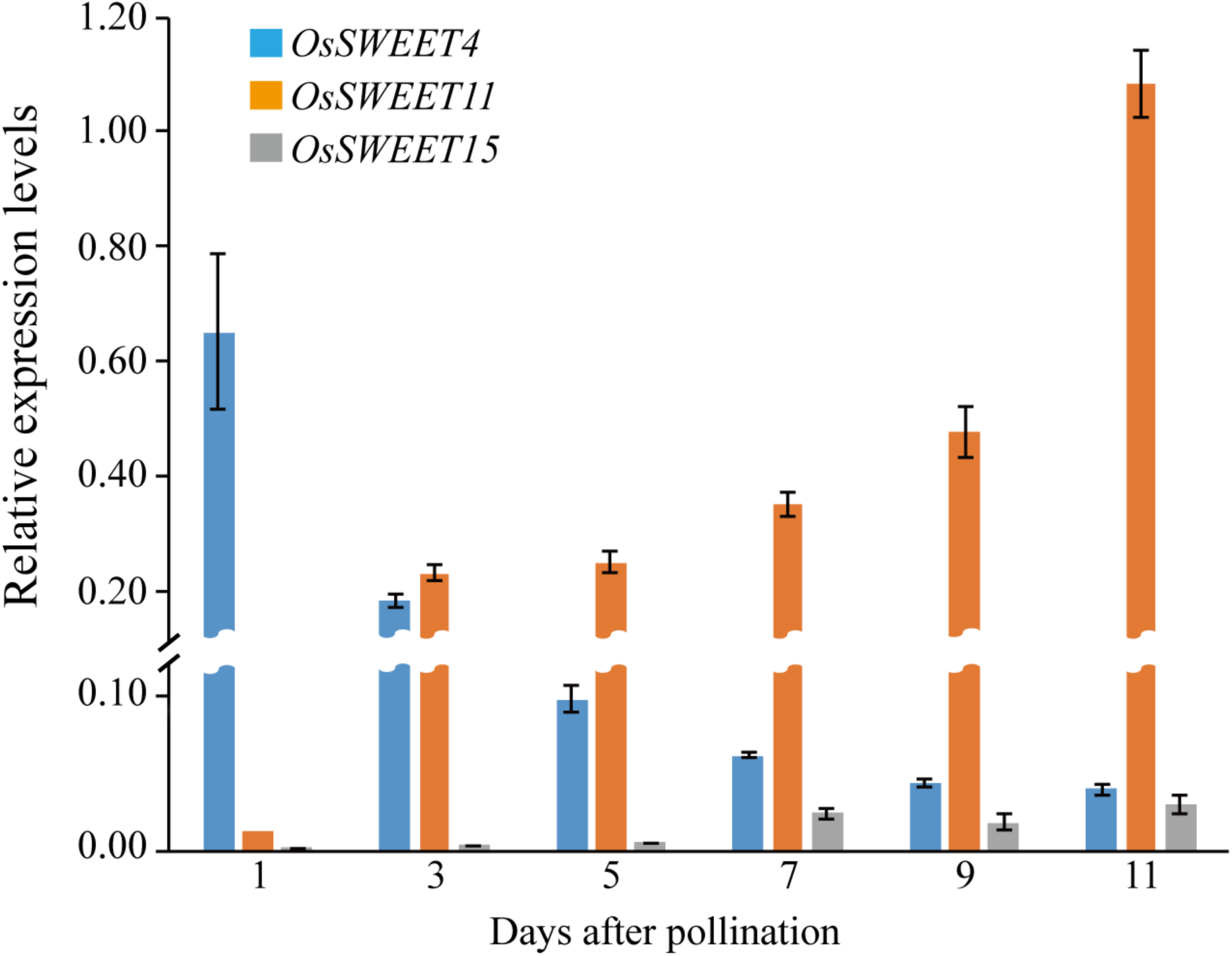
Relative expression of *OsSWEET4*, *OsSWEET11* and *OsSWEET15* during rice seed development. Expression was measured with quantitative RT-PCR (qRT-PCR) in wild-type greenhouse-grown seeds. Data was shown as mean ±s.e.m., n=3; expression levels were normalized to rice Ubiquitin1 levels.

### OsSWEET11 plays a key role in seed filling

OsSWEET11 had previously been shown to function as a plasma membrane sucrose transporter (Chen *et al.*, 2012). Since OsSWEET11 was by far the most highly expressed SWEET gene, we hypothesized that *knock-out* mutants might be affected in seed filling. Two independent *ossweet11* mutants, one carrying a single nucleotide deletion leading to a frameshift was created by CRISPR-Cas9 and a second TALEN-derived mutant carrying a 489 bp deletion were characterized phenotypically (Fig. S2). In the greenhouse, both mutants had incompletely filled seeds at maturity (Fig. 2a). Depending on the growth conditions, the phenotype was more or less severe (see for example Fig. 2 and Fig. S3). The effects became more severe in paddy field conditions (single field experiment in 2016; see also parallel study (Ma *et al.*, 2017). Moreover, panicle development of *ossweet11* was significantly delayed as apparent from panicles containing chlorophyll in the mutant still at 40 DAP (Fig. 2c,d). Maturity, i.e. loss of chlorophyll was completed in *ossweet11* mutants only much later (>60 DAP). As a result, mutants had a significantly reduced yield (both percentage of mature seeds after harvest and 1000- grain weight; Fig. S3a,b). Of note however, plant height, spikelet number and panicle length were similar as in wild-type also in paddy field conditions (Figs. 2c, S3c,d).

**Fig. 2.**
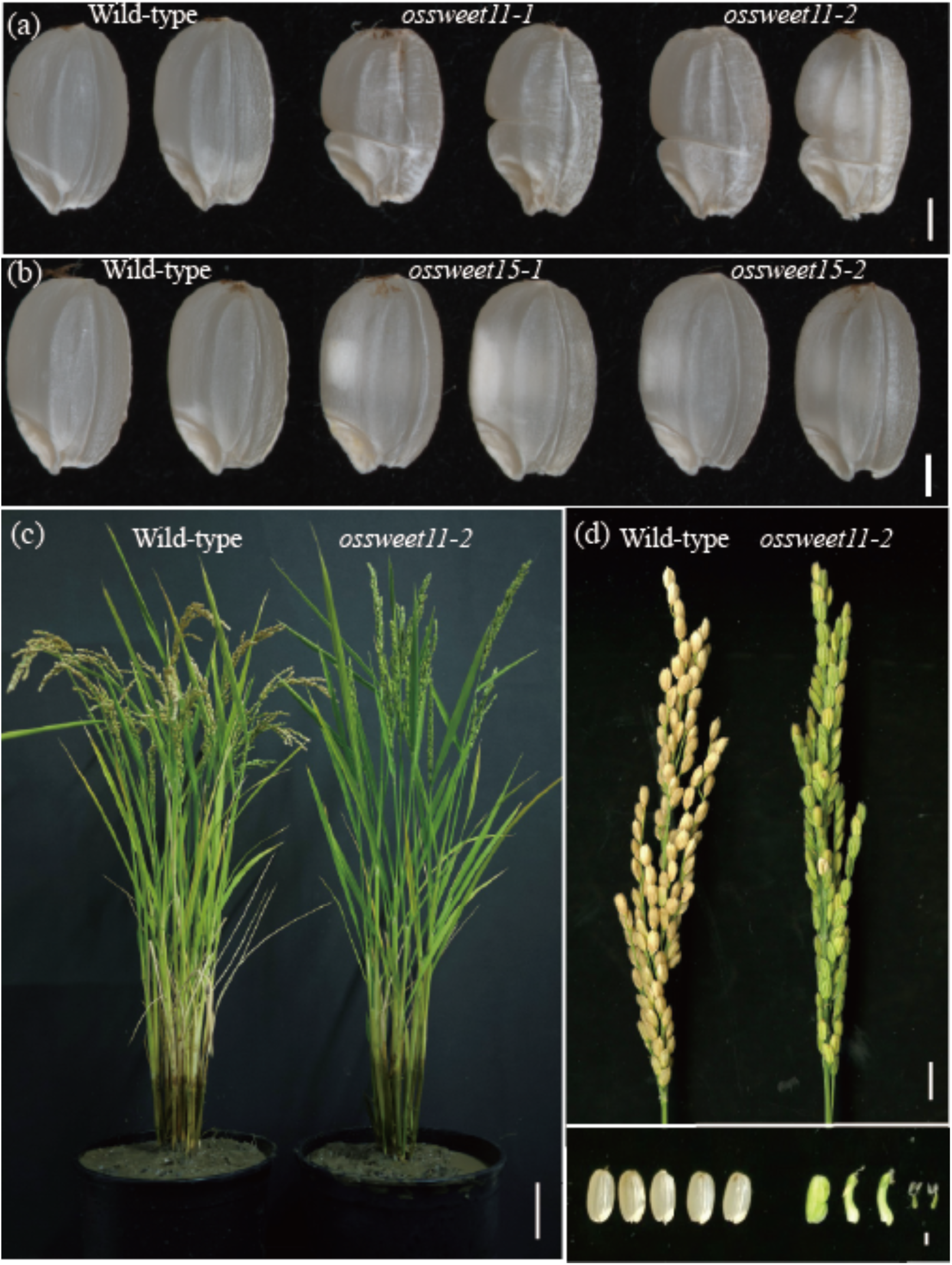
Phenotypes of wild-type, *ossweet11* and *ossweet15* mutants. **(a)** Mature caryopses phenotype of WT and *ossweet11* mutants (Stanford greenhouse). **(b)** Mature caryopses phenotype of WT and *ossweet15* mutants (Stanford greenhouse). **(c)** Plant phenotype of a mature *ossweet11-2 mutant* grown in a paddy filed and transferred to pot for photography (Stanford field, 40 DAP). **(d)** Phenotypes of panicle and grains for wild-type and the *ossweet11-2* mutant in paddy conditions (Stanford field, 40 DAP). Scale bars: 1 mm in (a and b), 10 cm in (c), 1 cm in upper panel and 1 mm in lower panel (d).

### OsSWEET11 accumulation in nucellar projection, nucellar epidermis and aleurone

Oparka had predicted symplasmic diffusion of sugar in the pericarp and apoplasmic transport at the nucellar epidermis-aleurone interface all around the endosperm, which contrasts the sucrose import patterns found in developing barely seeds, where the main import route was through the nucellar projection (Oparka & Gates, 1984; Melkus *et al.*, 2011). To determine whether OsSWEET11 exports sucrose at the nucellar projection or the nucellar epidermis/aleurone interface, we analyzed transgenic rice plants expressing translational GUS fusions containing a 2kb promoter fragment and the whole coding region including all introns. Crude histochemical GUS analysis of caryopses showed comparable GUS staining in seeds in 8 out of 18 independent transformants. Two independent lines were used for a more detailed analysis. In early stages (up to 3 DAP), we observe GUS activity in maternal tissues including the ovular vascular trace and the nucellus, possibly indicating a role in remobilization of carbohydrates during nucellar degradation (Fig. S4). At 5 DAP, GUS activity was detected in the ovular vascular trace, the nucellar projection, the nucellar epidermis surrounding the developing endosperm, the remaining nucellar proper and also in the aleurone layer of the endosperm (Fig. 3a-c). To our surprise, we find OsSWEET11 expression in nucellar projection as well as the nucellar epidermis, and in addition also in the outermost endosperm cell layer, the aleurone, providing a potential path for sucrose export out of the nucellar projection into the endosperm, and a parallel pathway for export from the circumferential nucellar epidermis and then subsequently a potential import via OsSWEET11 into the aleurone. A parallel study observed a similar expression pattern using a transcriptional GUS fusion (Ma *et al.*, 2017).

**Fig. 3.**
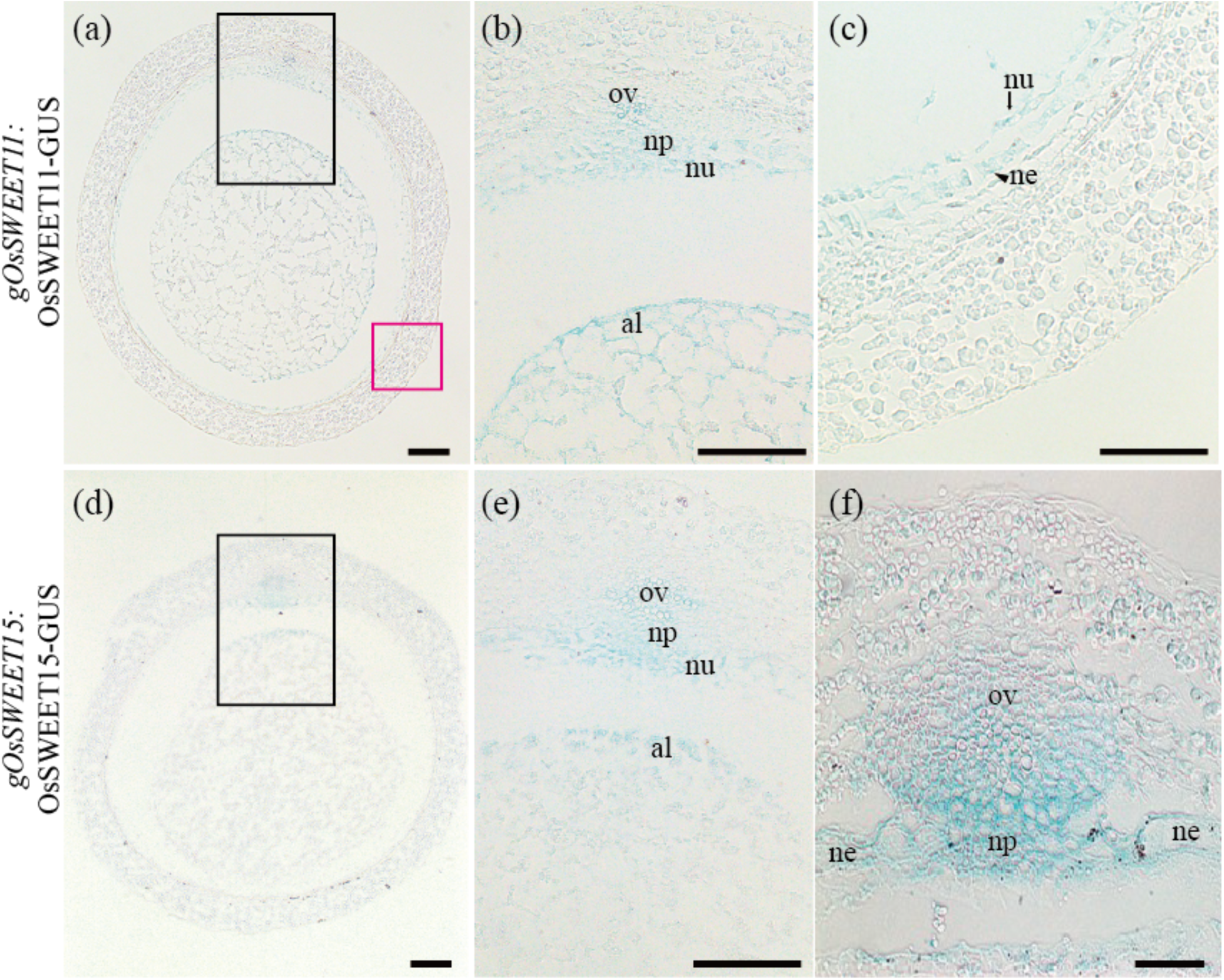
Tissue specific expression of OsSWEET11 and OsSWEET15 in rice grains.**(a)** Transverse section of *gOsSWEET11:*gOsSWEET11-GUS grain at 5 DAP. **(b)** Black boxed area in (a), showing GUS activity in the ovular vascular trace (ov), nucellus (nu), nucellar projection (np) and aleurone (al). **(c)** Red boxed area in (a), showing activity in the nu (black arrow) and ne, nucellar epidermis (black arrowhead). **(d)** OsSWEET15 GUS activity in grains at 5 DAP. **(e)** Black boxed area in (d), GUS activities detected in ov, np, nu and al. **(f)** OsSWEET15 GUS activity in developing grains was detected to the ov, np and ne at 9 DAP. Scale bars: 50 μm.

### **Potential compensation for *ossweet11* deficiency by other SWEETs**

The relatively weak phenotype of the *ossweet11* mutants may either indicate the existence of alternative genes and pathways, or could be due to compensation. Analysis of the expression levels of clade 3 SWEET genes in the *ossweet11* mutant showed that *OsSWEET13* mRNA levels were slightly increased, however the absolute levels were extremely low. The mRNA levels of *OsSWEET15* were about twofold higher in *ossweet11* seeds compared to wild-type (Fig. 4).Since depending on the experiment, *OsSWEET15* was expressed at only slightly lower levels compared to *OsSWEET11* and was furthermore candidate that may contribute to compensation in the mutant, we first tested whether it functions as a sucrose transporter, determined its expression pattern in developing caryopses and then analyzed knock-out mutants. As one may have predicted, OsSWEET15 also functioned as a sucrose transporter when co-expressed with a sucrose sensor in HEK293T cells (Fig. S5). Translational GUS fusions of the OsSWEET15 gene driven by their native promoter showed similar tissue specificity as compared to OsSWEET11 (13 independent lines): i.e. GUS activity was detected at early stages in the nucellus proper, and later in the ovular vascular trace, the nucellar projection and the aleurone (Fig. 3d,e). At 9 DAP, OsSWEET15 GUS activity was also detected in nucellar epidermis (Fig. 3f). The two SWEET transporters exhibit similar expression pattern in the developing seeds, especially ovular vascular and interface between the nucellar epidermis and the aleurone layer, assuming redundant roles during seed development. However, on their own, two independent *ossweet15 knock-out* mutants generated via CRISPR-Cas9 (frameshift mutations that prevent production of a functional OsSWEET15 protein, Fig. S2) did not show any detectable phenotypic differences compared to wild-type in 4 independent experiments (Fig. 2b).

**Fig. 4.**
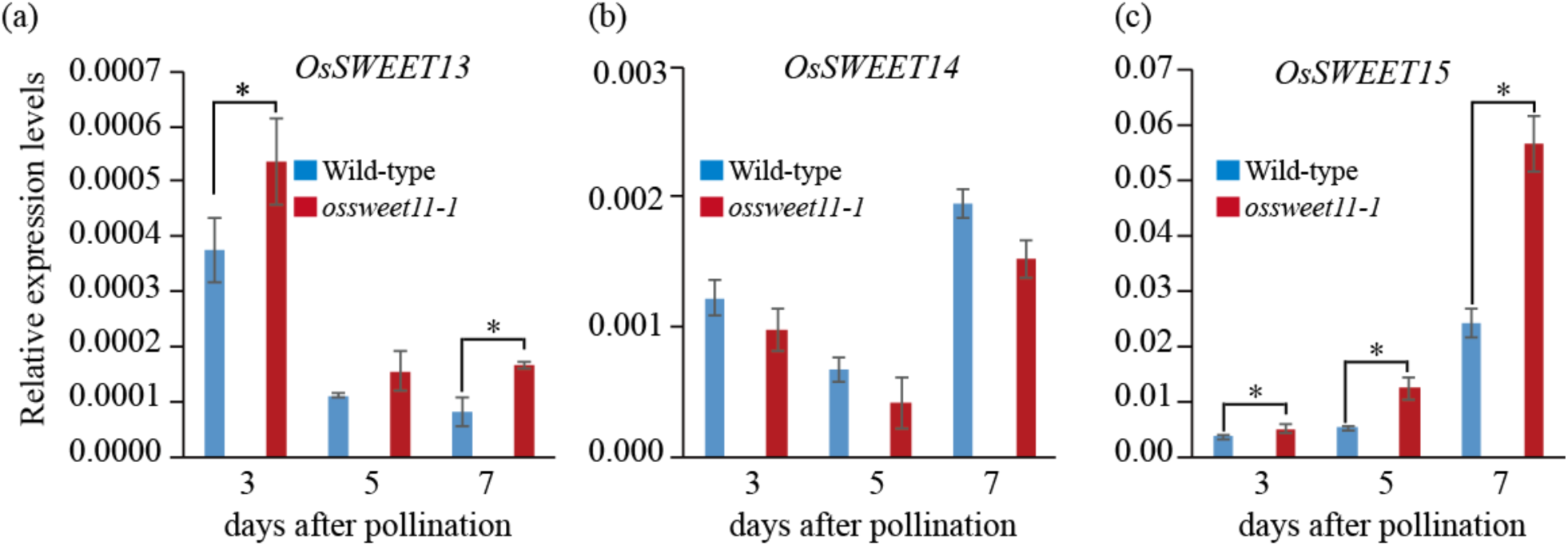
Expression levels of clade 3 SWEET genes in *ossweet11-1* mutant. Relative expression levels of *OsSWEET13* **(a)**, *OsSWEET14* **(b)** and *OsSWEET15* **(c)** was measured with quantitative RT-PCR (qRT-PCR) in wild-type and *sweet11*. Expression of *OsSWEET13* and *OsSWEET15* were increased in the *ossweet11-1* mutant compared to wild-type (* p<0.05). Data was shown as mean ± s.e.m., n=3; expression levels were normalized to rice *Ubiquitin1* levels. *OsSWEE12* transcripts were not detected.

### OsSWEET11 and 15 are essentially for seed filling

Since the seed filling of *ossweet11* mutants was only partially affected relative to *ossweet4* mutants (Sosso *et al.*, 2015), and OsSWEET15 appeared to be expressed in the same cell types to substantial levels, and even possibly compensates in part for OsSWEET11 deficiency in the mutant, we generated *ossweet11;15* double mutants for both alleles of the two loci. In greenhouse conditions both at ISU and Stanford, the double mutant phenotype was very severe, much more than the single *ossweet11* mutant (Fig. 5). The differences were even more severe in the ISU greenhouses, where *ossweet11*showed only a minor phenotype, whereas the caryopses of the double mutant were severely affected (Fig. S6). A detailed time series showed that phenotypic differences became apparent at ~5 DAP (Fig. 5a). Differences became much stronger at 7 DAP, a time point at which *ossweet11* mutants already started to develop a wrinkled grain morphology, while *ossweet11;15* was characterized by grains that were flattened with a smaller diameter (Fig. 5a,b). Sections through the grain showed that the mutants were endo-sperm-deficient and either had only remnants of the endosperm or lost the endosperm completely (Fig. 5c). Cytohistological analyses of resin-embedded sections showed that cellularization of the endosperm started at ~3 DAP in both wild-type and *ossweet11*, while cellularization was not observed in *ossweet11;15* mutant (Fig. S7). The cellularization of the endosperm was completed and nucellus were degenerated in the wild-type from 5 to 7 DAP (Fig. S7a). In the *ossweet11* mutant, endosperm was not fully cellularized and degradation of nucellus were delayed (Fig. S7b). In the *ossweet11;15* mutant, endosperm cells defected cellularization and nucellus remained until 7 DAP (Fig. S7c).

**Fig. 5.**
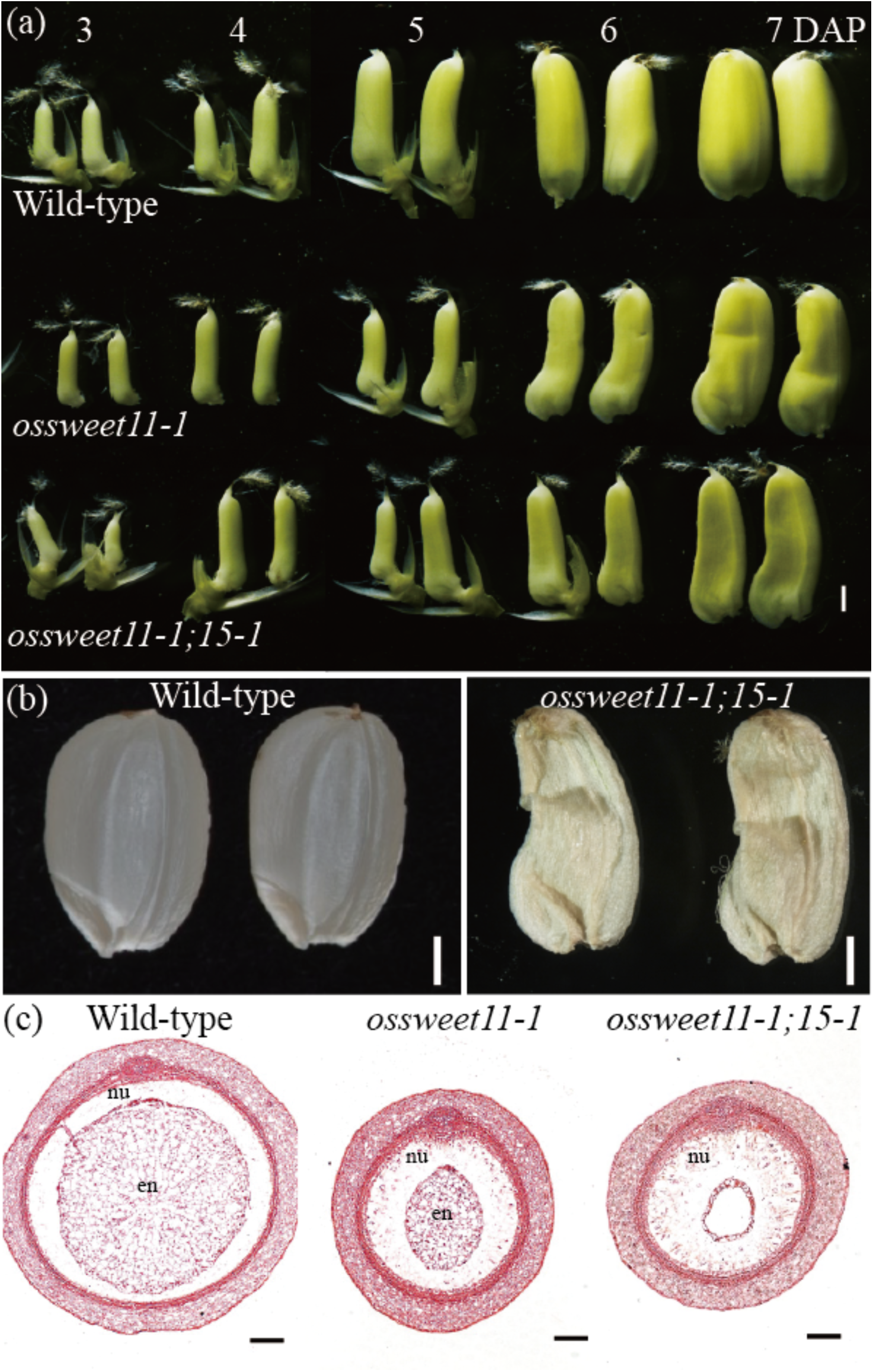
Endosperm deficiency phenotype of the *ossweet11;15* double mutant. **(a)** Morphological changes of wild-type, *ossweet11-1* and *ossweet11-1;15-1* from 3 DAP to 7 DAP (Stanford greenhouse). **(b)** Wild-type and *ossweet11-1;15-1* double mutant seeds at maturity (Stanford greenhouse). **(c)** Transverse sections of wild-type, *ossweet11-1* and *ossweet11-1;15-1* seeds at 5 DAP stained with Safranin O. nu, nucellus; en, endosperm. Scale bars: 1 mm in (a and b), 50 μm in (c).

### **Starch accumulation in the pericarp of *ossweet11;15* double mutants**

Based on the localization of OsSWEET11 and 15, we predicted that inhibition of the transporters would lead to starch accumulation in cells that export sucrose and cells outside the endosperm. In wild-type, starch is stored transiently in the pericarp until 7 - 9 DAP. Rapid starch degradation in the pericarp correlated with starch accumulation in the endosperm starting ~ 7 DAP (Wu *et al.*, 2016). We found starch in the endosperm, but only residual amounts in the pericarp of wild-type caryopses (Fig. 6a,d). By contrast, starch accumulated to high levels in the pericarp of the *ossweet11* mutant (Fig. 6b,e). In *ossweet11;15* double mutants, starch accumulated to even higher levels in the pericarp, while the endosperm did not show substantial starch levels (Fig. 6c,f). The accumulation of starch in the pericarp of *ossweet11* and *ossweet11;15* double mutants supports the critical roles of OsSWEET11 and OsSWEET15 in sugar translocation and mobilization towards the developing endosperm.

**Fig. 6.**
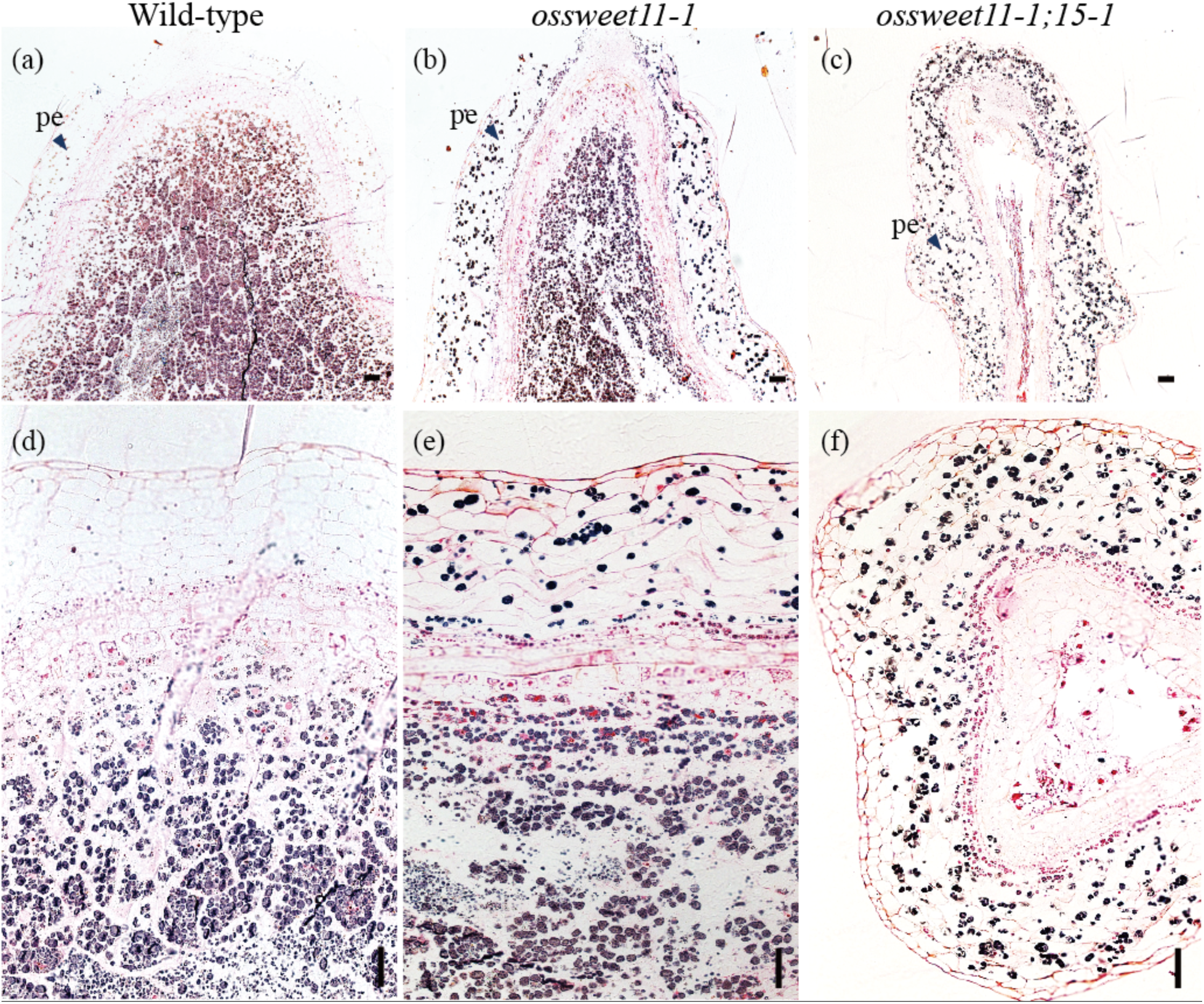
Accumulation of starch in the pericarp of *ossweet11-1* and *ossweet11-1;15-1* mutants. **(a-c)** Cross section of seeds stained with Lugol’ s iodine solution at 9 DPA. pe, pericarp. **(e-f)** Magnified image of (a-c). Scale bars: 50μm.

## Discussion

We draw five key findings from the results of a combination of analyses comprising gene expression, translational reporters to map tissue-specific protein accumulation, nuclease-induced knock out mutants: (i) OsSWEET11 and 15 are the most highly expressed sucrose transporting clade 3 SWEETs in the rice caryopsis, (ii) if we assume that they mark apoplasmic import routes, sucrose can enter both directly below the vein via the nucellar projection as well as the circumferential nucellar epidermis; (iii) they may not only play roles in cellular efflux at these two sites, but also be responsible for importing sucrose into the aleurone cells; (iv) OsSWEET11 and 15 are both contribute to seed filling with redundant roles, but since the single *ossweet15* mutant alone has no apparent phenotypic differences to wild-type, OsSWEET11 appears to function as the dominant transporter, consistent with the higher levels of mRNA; (v) *OsSWEET4*, since it is mainly expressed at early stages of development, may cooperate with the cell wall invertase cwINV2 (OsCIN2, GIF1) in hexose import in the vicinity of the dorsal vein, while OsSWEET11/15 are jointly contribute to seed filling at later stages. The findings made here for the *O. sativa* cv. Kitaake from OsSWEET11 are similar to those from a parallel study that used *O. sativa* cv. Nipponbare (Ma *et al.*, 2017). The main difference is that our lines showed a weaker phenotype when grown in greenhouse conditions relatively to the field experiments made with Nipponbare (Ma *et al.*, 2017).

### Pathways for seed filling

The tissue-specific expression of OsSWEET11 and 15 in parenchymatic cells of the vascular bundle, the nucellar projection, the nucellar epidermis and the aleurone layers in endosperm indicates specific role of OsSWEET11 and 15 in sucrose translocation into developing caryopses. Here we proposed possible model for sucrose transporting from ovular vascular bundle to endosperm (Fig. 7). There appear to be four locations that require sucrose transporters: **1. Parenchymatic cells in the vascular bundle**: a localization that may be expected based on the OsCIN2 localization which requires efflux of sucrose (and subsequent import of hexoses for providing sucrose as a substrate (Wang *et al.*, 2008). Of note, the *oscin2/gif1* cwINV mutant has a clearly distinct phenotypes with markedly more grain chalkiness and is thus not similar to *ossweet11* (Wang *et al.*, 2008). **2. Nucellar epidermis:** this is also an expected location that is compatible with Oparka’s work that indicates that the symplasmic pathway appears to be blocked, thus requiring transporters at the nucellar epidermis (Oparka & Gates, 1981a,b, 1982, 1984). **3. Aleurone:** an unexpected location which requires sucrose influx via plasma membrane transporters. The same localization of OsSWEET11 was also observed in a parallel study (Ma *et al.*, 2017). The authors found this unexpected, since it was a location that requires influx and is not expected to efflux sucrose. However, SWEETs appear to function as uniporters, thus the sucrose gradient simply determines the direction of flux. We had previously found that SWEET4 in maize and rice were likely key to the import of cwINV-derived hexoses in seeds (Sosso *et al.*, 2015). A sucrose gradient across the two cell types (nucellar epidermis and aleurone) driven by a high rate of delivery from the maternal side and rapid conversion in the endosperm would allow the use of the same transporters on both cell types. This situation is remotely similar as in the human intestine, where transcellular transport across the intestinal epithelia is mediated by GLUT2 on both the apical and basal membrane under conditions where the glucose concentrations in the lumen exceed those of the blood stream (Kellett *et al.*, 2008). In addition to the two SWEETs, *SUTs*, which are expressed in the aleurone, may contribute to secondary active sucrose import into the aleurone (Ishimaru *et al.*, 2001; Scofield *et al.*, 2002; Bai *et al.*, 2016). **4. Nucellar projection:** The presence of OsSWEET11 and 15 in the cells of the nucellar projection may appear as the most surprising site, since plasma membrane sucrose transport is not in line with radiotracer import studies, which indicated that in rice, the import of sugars occurs exclusively via the nucellar epidermis-aleurone pathway (Oparka & Gates, 1981b).However, others have suggested that the nucellar projection may help to transporting sugars to the developing endosperm also in rice (Krishnan & Dayanandan, 2003). Importantly, the nucellar projection pathway appears to be the main pathway for sugar import in barely caryopses as shown by magnetic resonance imaging (MRI)(Melkus *et al.*, 2011). Of note, we are aware that by contrast to Oparka’s radiotracer studies, we do not measure actual translocation of assimilates, but rather the presence of a protein, and we do not know whether the two SWEETs are active at the plasma membrane of these cells. Nevertheless we suggest that it may be useful to reassess sugar entry pathways, for example by MRI at different stages and in different varieties.One possible difference could be that Oparka used an *indica* rice variety (IR 2153-338-3), while the *japonica* variety Kitaake was used for all experiments shown here; Nipponbare was used by the parallel study that localized a transcriptional GUS fusion of the OsSWEET11 promoter to the same cells as the translational fusions in our work (Ma *et al.*, 2017). Alternatively, these pathways may be used at different stages of development. Notably, the three Arabidopsis transporters SWEET11, 12 and 15 which play critical roles for sucrose efflux from the seed coat also showed very complex changes in cellular expression during seed development (Chen *et al.*, 2015b).

**Fig. 7.**
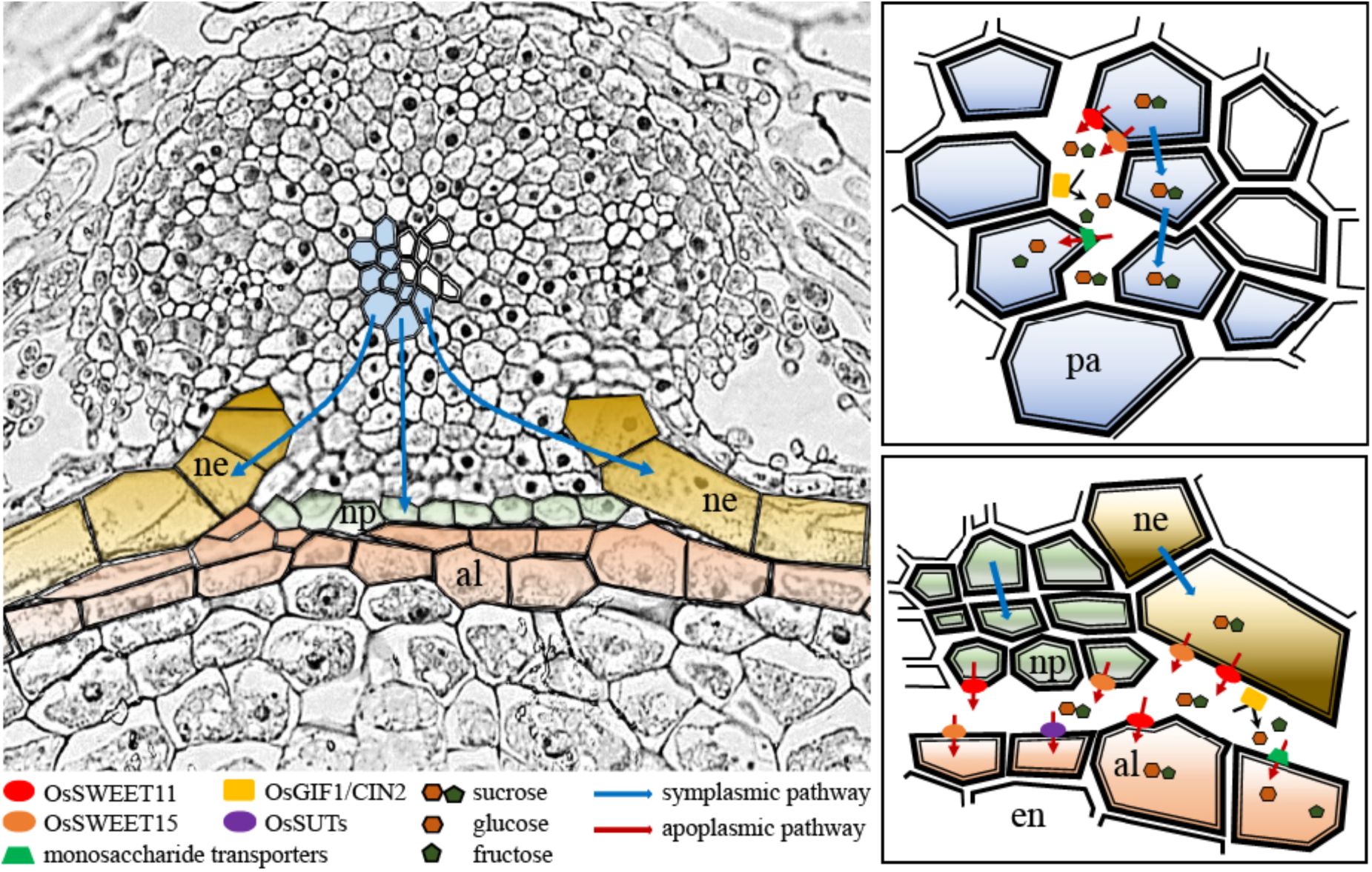
Proposed model for sugar unloading in rice caryopsis. Possible apoplasmic transport routes as indicated by SWEET sucrose transporter localization.The model was made based morphological observations from this study and previous studies (Oparka & Gates, 1981a,b; Wu *et al.*, 2016). Sucrose may move from the phloem to parenchyma cells in the ovular vascular bundle and then to nucellar projection and the nucellar epidermis through symplasmic pathway via plasmadesmata **(a)**. We surmise that OsSWEET11 and OsSWEET15 mediates sucrose export from xylem parenchyma cells into the apoplasmic space **(b)**. In addition, OsSWEET11 and OsSWEEET15 may be involved in sucrose export out of cells at the nucellar projection and the nucellar epidermis to apoplasm. followed by import into the aleurone in endosperm **(c)**. Transfer across the nucellar epidermis/aleurone would require a sucrose gradient across both cell types. al, aleurone; ne, nucella epidermis; np, nucellar projection; pa, parenchyma.

### Starch in the pericarp as a transient buffer

In rice caryopsis development, large amount of starch grains accumulated in pericarp at 6 DAP, followed by those starch grains degraded from 7 to 9 DAP (Wu *et al.*, 2016). This type of starch accumulation and degradation has been observed in pericarp of barely and wheat (Radchuk *et al.*, 2009; Xiong *et al.*, 2013). Starch accumulation occurred in the *ossweet11* and *ossweet11;15* of pericarp at 9 DAP (Fig. 6), suggesting the presence of one route for delivery of sucrose in the pericarp.

Ma et al. (Ma *et al.*, 2017) also localized transcriptional reporter fusions of OsSWEET11 to the pigment strand close to the main vascular trace and found a severe seed filling phenotype when analyzing plants grown in the field. In our greenhouse experiments, the phenotypic effect of *ossweet11* mutations was a lot less severe, in some cases even marginal, intimating a strong effect of the growth conditions, possibly light and nutrition on the phenotype. Our work indicates, based on the similarity in steady state RNA levels, timing of mRNA accumulation, tissue specificity and the combined effect observed in double knock out mutants that OsSWEET15 can compensate for OsSWEET11 deficiency.

### Developmental control of seed filling

Rice caryopsis occurred dynamical changes in cell expansion, cell wall thickening, and starch grain accumulation (Wu *et al.*, 2016). Expression levels of *OsSWEET11* and *OsSWEET15* are gradually increased during caryopsis development. In contrast *OsSWEET4*, which is expressed mainly at the base of the caryopsis, and mutation strong seed filling defect, however timing very different, very high early and then declining (Fig.1). Early seed development, imported sucrose released from maternal tissue is cleaved by an extracellular invertase. Similarly with *OsSWEET4*, *OsCIN1* and *OsCIN2* are mainly expressed early stage (Hirose *et al.*, 2002; Cho *et al.*, 2005). Since OsSWEET4 belongs to clade 1 SWEET and shown as a glucose transporter (Sosso *et al.*, 2015), role of clade 1 glucose transporter might be important for early seed development, but clade 3 SWEETs might be important for late seed development.

### Relevance for pathogen susceptibility

The finding that OsSWEET11 and 15 play important roles in seed filling is also relevant in the context of the fact that OsSWEET11 serves as a blight susceptibility gene (Yang *et al.*, 2006; Yuan *et al.*, 2010; Antony *et al.*, 2010; Chen *et al.*, 2010). Ectopic expression of *OsSWEET11* is activated by pathovar-specific effectors of the blight pathogen *Xanthomonas oryzae pv oryzae*, and mutations in the effector binding sites in the *OsSWEET11* promoter lead to resistance to Xoo (Yang *et al.*, 2006; Yuan *et al.*, 2010; Antony *et al.*, 2010). Therefore, it will be important to ensure that engineering of the OsSWEET11 promoter in resistant lines retains proper OsSWEET11 expression in seeds to ensure that resistant lines do not carry a yield penalty. This goal appears feasible since apparently mutants (*xa13*) that are used by breeders do not show yield deficiencies (Laha *et al.*, 2016).

## Conclusions

The analysis of SWEET gene expression in rice caryopses together with the characterization of knockout mutants in the most highly expressed clade 3 SWEETs 11 and 15 demonstrates that OsSWEET11 and 15 play central roles in seed filling. The cellular expression patterns of OsSWEET11 and 15 indicate that there may be multiple apoplasmic pathways for sucrose entry into the endosperm. A careful analysis of the timing and localization of other sugar transporters of the SWEET, SUT and MST families as well as the cell wall invertases, and an analysis of sucrose import by MRI in a variety of rice cultivars will help to further delineate the sugar import pathways and hopefully contribute to the knowledgebase for engineering improve yield potential in rice.

## Acknowledgments

We thank Davide Sosso for helpful discussions. This research was supported by grants from the National Science Foundation research (IOS-1258103) to WF and BY and the Department of Energy (DE-FG02- 04ER15542) to WF.

## Author contributions

JY, JSE, BY and WF conceived and designed the experiments; JY, JSE and DL performed the experiments and collected the data, executed the data analyses, and rendered the figures; and all authors contributed to the interpretation of the results, wrote and revised earlier drafts @ they approve the final version of this manuscript and agree to be held accountable for the content.

